# Development of a mouse model for postresectional liver failure

**DOI:** 10.1101/2022.11.10.515992

**Authors:** Kim M.C. van Mierlo, Cathy Van Himbeeck, Valérie Lebrun, Peter L.M. Jansen, Cornelius H.C. Dejong, Isabelle A. Leclercq, Steven W.M. Olde Damink, Frank G. Schaap

## Abstract

**Background:** Postresectional liver failure (PLF) is a dreaded complication after extended liver resection. Post-operative hyperbilirubinemia suggests that impaired hepatobiliary transport with intrahepatic accumulation of harmful cholephiles plays an etiological role. Bile salts serve dual roles as signaling molecules engaged in liver regeneration after partial hepatectomy (PH) and biological detergents.

**Aim:** In this study we tested the hypothesis that excessive accumulation of bile salts in the regenerating liver results in PLF.

**Methods:** Twelve weeks old male C57BL6/J mice were subjected to 70% PH and post-operatively challenged with a diet supplemented with cholic acid (CA, 0.5 or 1.0%; n=5-6 per group) or a control diet. After 48 hours mice were sacrificed, and liver injury, secretory function, and regenerative indices were assessed.

**Results:** Mice fed a 1.0% CA diet displayed more pronounced weight loss following PH and had a deranged post-operative glucose course. Liver injury (aminotransferase elevations) and impaired hepatobiliary transport function (hyperbilirubinemia) were apparent in the group fed a 1.0% CA diet, but not in animals fed a 0.5% CA diet. No differences in liver mass recovery were observed among groups. However, the percentage of hepatocytes staining positive for the proliferation marker Ki-67 were reduced in mice receiving a 1.0% CA diet relative to animals fed a 0.5% CA diet. PH-induced expression of key factors involved in cell cycle progression (*e*.*g. Foxm1b, Cdc25b*) was abrogated in the 1.0% CA group.

**Conclusion:** A postresectional challenge with a 1.0% CA diet induces signs of liver injury and defective liver regeneration. A longer duration of the dietary challenge and/or secondary hits may further improve the model. Once validated, it can be used to evaluate pharmaceutical strategies to prevent or treat PLF.

## 1. Introduction

Annually, approximately 7.000 patients develop primary liver cancer or liver metastases in the Netherlands alone. Liver resection is the preferred curative treatment for liver malignancies. The feasibility of this procedure is largely dependent on tumor mass extension and localization, resulting in only 15-20% of patients with liver cancer qualifying for resection [1]. Although liver resection is a relatively safe intervention, it is still associated with a mortality rate as high as 2-5% in patients with colorectal liver metastasis and up to 10% in patients with perihilar cholangiocarcinoma [2]. Up to 75% of this mortality can be attributed to postresectional liver failure (PLF), where the capacity of the remnant liver to regenerate and simultaneously exert its normal functions, is hampered due to an inadequate quantity and quality of the residual liver mass [1,3].

Post-operative elevation of bilirubin, used in all current clinical definitions of PLF, suggests that impairment of hepatobiliary transporters may underlie development of PLF [3]. Moreover, after liver resection, the remnant liver faces a relative overload of bile salts (BS) because the original BS pool recirculates through a smaller liver remnant that apparently has insufficient spare capacity to properly handle this increment. This results in increased systemic spill-over and elevation of circulating bile salts [4]. BS are known to bind to and - if agonistic-activate several nuclear receptors (NRs) including farnesoid X receptor (FXR or NR1H4) and pregnane X receptor (PXR). The ligand-activated transcription factor FXR is highly expressed in the small intestine and liver, but also in the adrenal glands and kidneys [5]. Upon activation, transcriptional induction of target genes of FXR such as the canalicular bile salt export pump (BSEP) maintain hepatic BS homeostasis, thus, limiting BS toxicity and subsequent liver injury. In addition, FXR is responsible for feedback inhibition of hepatic BS synthesis via upregulation of ileal fibroblast growth factor 19 (FGF19) and hepatic small heterodimer partner (SHP), which target expression of the rate-limiting enzyme (cholesterol 7a-hydroxylase, encoded by the *CYP7A1* gene) through different mechanisms [5,6]. The enterokine FGF19 (Fgf15 in rodents) reaches the liver via the portal circulation, and acts via the FGFR4/βKlotho complex on the cell surface of hepatocytes.

Mouse studies have shown that *Fgf15* KO mice have increased levels of hepatic *Cyp7a1*, both at mRNA as well as protein level, with a parallel increase in enzyme activity [7]. The role of Fgf15 in liver regeneration was demonstrated by Uriarte et al. (2013), who showed the importance of Fgf15 in maintaining BS homeostasis and preventing liver damage and mortality after partial hepatectomy (PH) [8]. Moreover, this study also revealed that Fgf15 mediates spontaneous liver growth, *viz*. in the absence of a surgical trigger, by a diet containing the bile salt cholic acid (CA).

Systemic and intrahepatic accumulation of bile salts are considered a causative factor in acute liver failure (ALF)[1], as observed in mice models of extended hepatectomy (85% liver resection) [9]. In these models, the relative overload of BS after hepatectomy caused hepatocellular injury and resulted in impaired liver regeneration and increased mortality [9,10]. We hypothesized that excessive accumulation of BS in the regenerating liver is the actual culprit in PLF. To test this hypothesis, we aimed to develop a mouse model of PLF by inducing BS overload in the regenerating liver.

## 2. Materials and methods

### 2.1 Animal handling and partial hepatectomy procedure

Male C57BL6/J mice (n=18, 11 weeks old) were obtained from Harlan Laboratories (Horst, the Netherlands) and housed in the animal facility of the Université catholique de Louvain (UCL; Brussels, Belgium). The animals were kept under controlled conditions with exposure to a 12-h light/12-h dark cycle and a constant temperature of 20-22°C. Animal experiments and care were conducted in accordance with European regulations and FELASA guidelines for humane care for laboratory animals provided by UCL. Postoperative welfare was assessed with a welfare scoring sheet (**Table S1**). The study protocol was approved by the university ethics committee (ref nr 2012/UCL/MD/026). All mice were fed standard chow (Global diet 2016; Harlan Laboratories, Madison, USA) for one week, after which they were subjected to 70% PH (T=0). The procedure was performed essentially according to the protocol of Mitchell and Willenbring [11], whereby in this study only a small abdominal incision was made and the gallbladder was removed simultaneously. After the weight of excised median and lateral lobes (‘anterior/resected lobes’, ∼70% of liver mass) was recorded, they were fixed in 4% formalin or snap-frozen in liquid nitrogen and stored at −80°C until analyses. After resection, mice (n=5-6 per group) were fed the same diet supplemented with 0.0%, 0.5% or 1.0% CA (Custom diet, Harlan Laboratories). Mice were sacrificed by exsanguination 48 hrs after 70% PH. One mouse (0.0% CA diet group) was excluded from the analyses based on elevated liver enzymes, deviant gene expression levels and elevated liver bile salt levels, likely resulting from a technical failure of the surgical procedure.

At the time of sacrifice, mice were anesthetized via intraperitoneal injection with ketamine/xylazine. Ten minutes later, the abdominal cavity was opened via a midline incision and blood was drawn by portal vein puncture, kept on ice, and serum was prepared and stored at −80°C until further use. After harvesting and weighing the liver (‘posterior lobes’), the ileum (distal ⅓ part of the small intestine) was also harvested and both tissues were fixed in 4% formalin, or snap-frozen in liquid nitrogen and stored at −80°C until analysis.

The rate of liver mass recovery was estimated using the following formula:

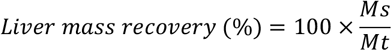

where M_s_ is the liver weight at sacrifice, and M_t_ is the total liver mass before PH (estimated by dividing the mass of resected segments by 0.7).

### 2.2 Biochemical analyses

Body weight (g) of the animals was monitored daily during weekdays at set time points. Blood glucose levels (mg/dL) were measured in blood drawn from the lateral tail vein (Accu-chek Aviva, Roche diagnostics, Mannheim, Germany) every 4 hours after PH, with a final measurement 2 hours prior to sacrifice. Serum aminotransferases (ALT, AST) and bilirubin were determined 48 hours after PH via automated procedures (Department of Bio-Medical Chemistry and Clinical Biology, St Luc University Hospital, Brussels, Belgium).

### 2.3 Immunohistochemistry

Hepatocyte proliferation was assessed via Ki-67 immunohistochemical staining. Firstly, liver tissue was fixed in 4% formalin, dehydrated in graded ethanol and embedded in paraffin to allow the creation of serial tissue sections of 4 μm thickness. Mouse monoclonal antibody against Ki-67 (dilution 1:50; Code No. M7249, Dako, Glostrup, Denmark) was used. Anti-mouse Envision system (Dako) was used for secondary detection. For visualization, the 3,3’-Diaminobenzidine (DAB) Substrate-Chromogen System (Dako) was used. Nuclei were counterstained with haematoxylin. The proliferative index (%) was measured by dividing the amount of Ki-67 positive hepatocyte nuclei by the total number of hepatocyte nuclei in four high-power (40x) fields.

### 2.4 Total bile salt assay

A 5% liver homogenate was made by homogenizing ca. 50 mg of tissue in 1 mL of 75% ethanol by means of a Mini-Beadbeater (Biospec Products, Bartlesville, USA). Thereafter, BS were extracted with 75% ethanol by incubating the homogenates for 2 hours at 50°C [12]. Supernatant was collected after centrifugation (10 min., 20620*g* at 4°C). Total BS in liver extract or serum was quantified with an enzymatic assay according to the manufacturer’s protocol (Total bile acid assay, Diazyme Laboratories, Dresden, Germany). The amount of BS present in liver extracts was normalized to wet liver mass.

### 2.5 Assessment of gene expression

Expression levels of genes involved in BS homeostasis, cell cycle regulation and proliferation, were measured in liver and ileum samples via RT-qPCR. To study the effect of PH on gene expression *per se*, non-regenerated liver samples (resected lobe, T=0h) were included. RNA was isolated from a resected liver lobe (T=0), the regenerating lobe at sacrifice (T=48h), as well as ileum tissue (T=48h), with TRI reagent solution according to the manufacturer’s protocol (Sigma-Aldrich, St. Louis, Missouri). The concentration and purity of RNA were determined by measuring absorbance with a NanoDrop 1000A spectrophotometer (Thermo Scientific). After isolation, RNA was treated with DNAse (Promega, Madison, Wisconsin) to degrade residual genomic DNA, efficiency of DNAse treatment was verified by PCR. Next, 750 ng total RNA was reverse transcribed to form cDNA according to the manufacturer’s protocol (SensiFAST™ cDNA Synthesis Kit, Bioline, Luckenwalde, Germany). Real-time PCR analyses were performed on a MyiQ Single-Color Real-Time PCR detection system (Bio-Rad, Veenendaal, the Netherlands), employing SYBR Green chemistry (SensiMix™ SYBR® & Fluorescein Kit, Bioline). PCR reactions contained 2.0 μL diluted cDNA sample (corresponding to 7.5 ng total RNA) in a total volume of 10 μL. qPCR data was analyzed using LinReg software [13]. Results were normalized using *Rplp0/36b4* as a reference gene. Data are expressed as fold change relative to the median expression in the control group. Sequences of primer pairs used for RT-qPCR analyses are provided in **Table S2**.

### 2.6 Statistical analysis

The experimental results were statistically analysed using IBM SPSS statistics version 22. Friedman test with Dunn’s multiple comparisons testing was used to analyze the evolution of body weight and glycaemia during the study period for all groups (n=5-6 per dose group). Pre-operative (day −5 until T=0) and post-operative (from T=0 until sacrifice) trajectories were analyzed separately. Between-group comparison of glucose levels at each time point, was performed using the Kruskal-Wallis test. The effect of PH (resected lobes at T=0 [n=9] vs. T=48 regenerating lobes of 0.0% CA group [n=5]) was analyzed via a Mann-Whitney U test. *P*-values ≤ 0.05 were considered statistically significant. Differences between the three dose groups (n=5-6 per group) were analyzed by means of a Kruskal-Wallis test. If the Kruskal-Wallis test showed significance (*P*≤0.05), post-hoc Mann-Whitney U tests were performed between all groups with Bonferroni-Holm correction for multiple comparisons. For visual purposes, biochemical data in graphs is depicted as mean ± SEM. All experimental data on gene expression is graphically presented as median with interquartile range.

## 3. Results

### 3.1 Effect of a CA diet on evolution of body weight, glycaemia, and liver regeneration

During the pre-operative course in which mice were fed normal chow, significant weight gain was observed in time (*P*_time_ <0.001-0.0370), **Fig. 1A**). PH resulted in a decrease in body weight in all groups (*P*_time_<0.001). At postoperative day one, the highest dose group suffered from more pronounced weight loss compared to the 0.0% and 0.5% CA diet groups (*P*=0.016 resp. *P*=0.009).

**Figure 1.**
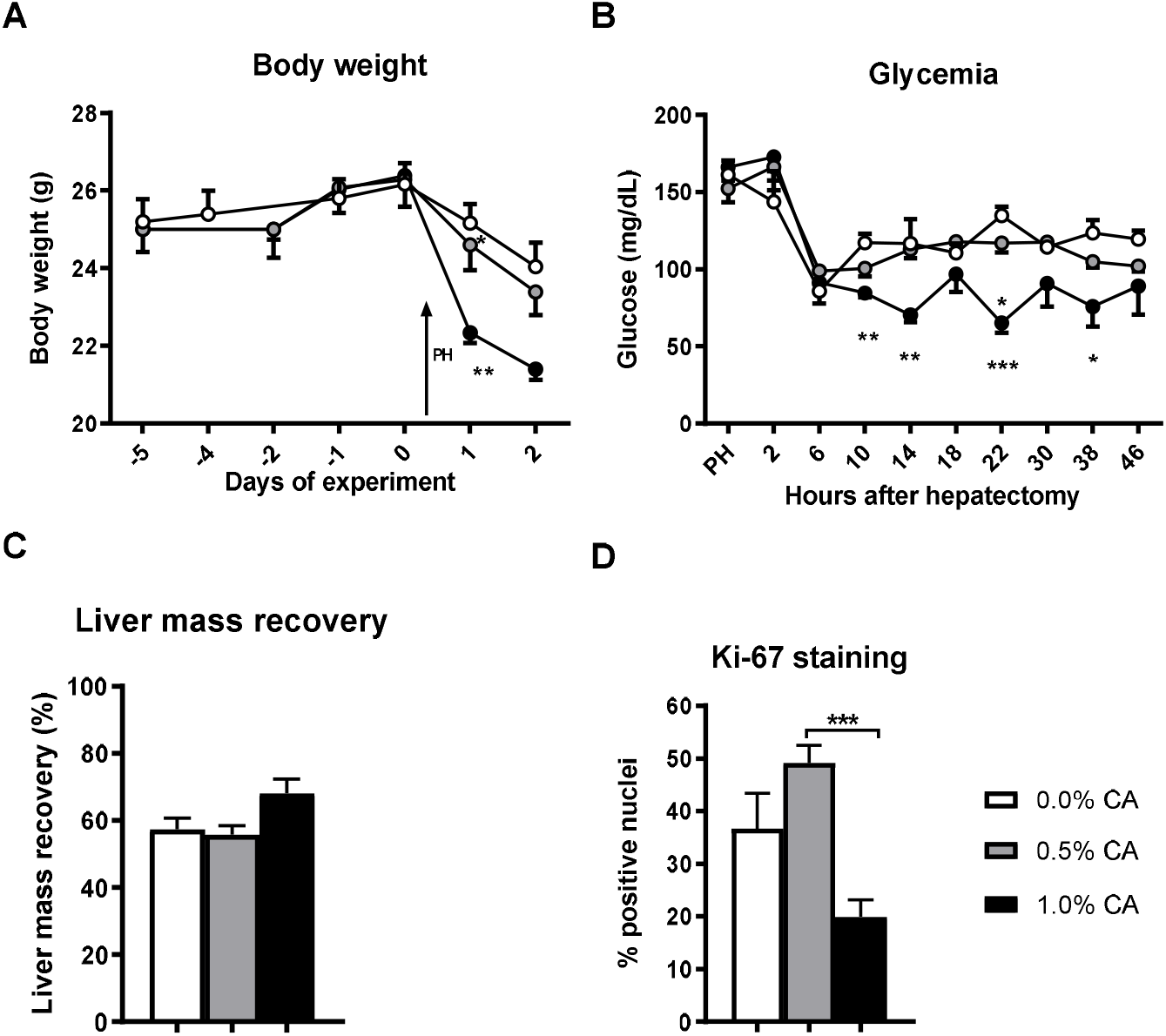
Metabolic derangement and abrogated hepatocyte proliferation in hepatectomized mice fed a diet containing 1.0% cholic acid. Mice (n=5-6 per group) were subjected to 70% partial hepatectomy (PH) and post-operatively challenged with a diet containing 0.0%, 0.5% or 1.0% cholic acid (CA). Mice were sacrificed at 48 hrs after PH. Experimental courses of body weight (**A**) and blood glucose (**B**). Data are presented as mean with SEM. Regeneration after PH was assessed by recovery of liver mass (**C**) and immunohistochemical analysis of hepatocyte proliferation (**D**). Data are presented as mean with SEM. \**P*<0.05, \*\**P*<0.01,\*\*\**P*<0.005.

Before PH, glucose levels were not different between the three groups (**Fig. 1B**). PH induced a transient decrease in glucose levels in all groups (*P*_time_=0.005 for 0.0% CA, *P*_time_<0.001 for 0.5 and 1.0% CA), with mice fed a 1.0% CA diet showing an overall deviating course compared to the two other groups. Serum glucose was significantly reduced in mice receiving diet containing 1.0% CA, at T=10, T=14, T=22 and T=38h (**Fig. 1B**). No differences in liver mass recovery (based on wet liver mass) were observed between the three groups (**Fig. 1C**). However, mice fed a 1.0% CA diet were found to have a lower percentage of Ki-67^+^ hepatocytes in comparison with mice fed a 0.5% CA diet (49% vs. 20%, *P*=0.002, **Fig. 1D**).

### 3.2 Effect of dietary cholic acid on markers for liver injury and secretory function

Mice fed a 1.0% CA diet had significantly increased levels of alanine aminotransferase (ALT) and aspartate aminotransferase (AST) in comparison to the 0.0% CA diet group (both *P*=0.004, **Fig. 2AB**). Moreover, serum AST was increased in mice fed a 0.5% diet compared to the 0.0% CA diet group. Regarding total and direct bilirubin levels, mice fed a 1.0% CA diet showed higher levels than mice fed a 0.5% CA diet (*P*=0.002 resp. *P*=0.007; **Fig. 2CD**), indicating an impaired secretory function of the liver in the highest dose group. Moreover, mice in the 1.0% CA group did ‘clinically’ worse than the other groups as observed by reduced physical activity, hunched posture, squinted eyes and piloerection. At sacrifice, their livers appeared yellow and ascites was observed (**Fig. S1**).

**Figure 2.**
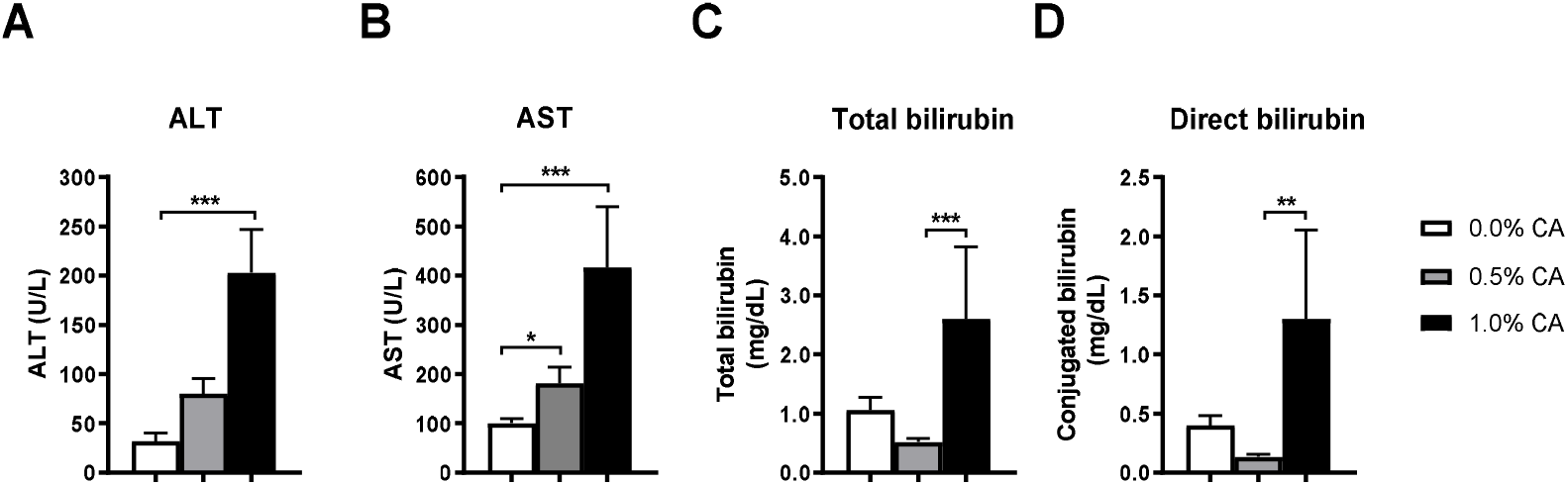
Elevated liver enzymes and hyperbilirubinemia in hepatectomized mice fed diet containing 1.0% cholic acid. Mice were subjected to 70% partial hepatectomy (PH) and post-operatively challenged with diet containing 0.0%, 0.5% or 1.0% cholic acid (CA, n=5-6 per group). Mice were sacrificed at 48 hrs after PH. Liver injury after hepatectomy was assessed by serum liver enzymes (**A**,**B**) and secretory function (**C**,**D**). Data are presented as mean with SEM. \**P*<0.05, \*\**P*<0.01,\*\*\**P*<0.005. CA, cholic acid; ALT, alanine aminotransferase; AST, aspartate aminotransferase.

### 3.3 Influence of dietary cholic acid on hepatic and serum bile salt content

After PH, hepatic BS content dropped significantly (T=0 vs. T=48 hrs in the 0.0% CA group, *P*=0.009, **Fig. 3A**). After PH, mice fed a 0.5% and 1.0% CA diet both showed significantly higher hepatic BS levels compared to a 0.0% CA diet (*P*=0.004 resp. *P*=0.017; **Fig. 3A**). Serum BS levels were increased in the mice fed a 0.5% CA diet in comparison to the control group and mice fed a 1.0% CA diet. (*P*=0.004 resp. *P*=0.015, **Fig. 3B**).

**Figure 3.**
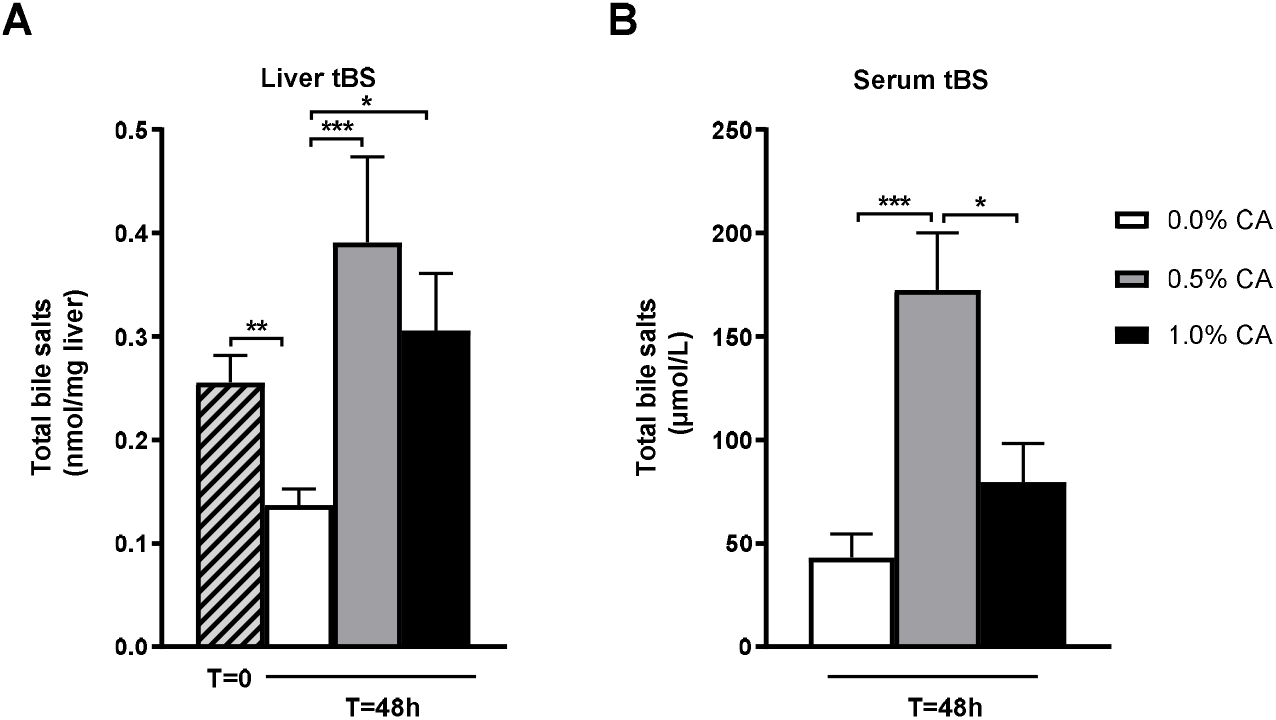
Disturbed bile salt homeostasis in the regenerating liver upon feeding a cholic acid-containing diet. Mice were subjected to 70% partial hepatectomy (PH) and post-operatively challenged with diet containing 0.0%, 0.5% or 1.0% cholic acid (CA, n=5-6 per group). Mice were sacrificed at 48 hrs after PH. Total bile salt levels were measured in liver before and after hepatectomy (**A**) and in serum after hepatectomy (**B**). Data are presented as mean with SEM. \**P*<0.05, \*\**P*<0.01,\*\*\**P*<0.005. CA, cholic acid; tBS, total bile salts.

#### 3.3.1 Effect of a post-operative cholic acid diet on bile salt synthesis and regulation

Maintenance of bile salt homeostasis in the remnant liver is a prerequisite for normal progression of liver regeneration after PH. Notably, hepatic BS accumulation was accompanied by reduced hepatocyte proliferation in mice receiving diet with 1.0% CA. Expression of genes engaged in different aspects of bile salt homeostasis was determined to investigate this further.

Expression levels of genes involved in BS synthesis and regulation thereof, were measured in liver and ileum. The regulatory genes include ileal *Fgf15* and hepatic *Fxr* and its direct target gene *Shp*, while genes engaged in BS synthesis include *Cyp7a1, Cyp8b1* and *Cyp7b1* (liver). PH induced a downregulation in hepatic gene expression of *Fxr* and *Shp* (*P*<0.001 resp. *P*=0.029) in the 0.0% CA diet group (**Fig. 4AB**). No differences in expression of *Fxr* and *Shp* in the different diet groups were observed after PH. After PH, gene expression of ileal *Fgf15* was elevated 16.5-fold in mice fed a 0.5% CA diet in comparison with a 0.0% CA diet (*P*=0.011) (**Fig. 4C**). However, no significant difference was found in comparison with the 1.0% CA group (*P*=0.628).

**Figure 4.**
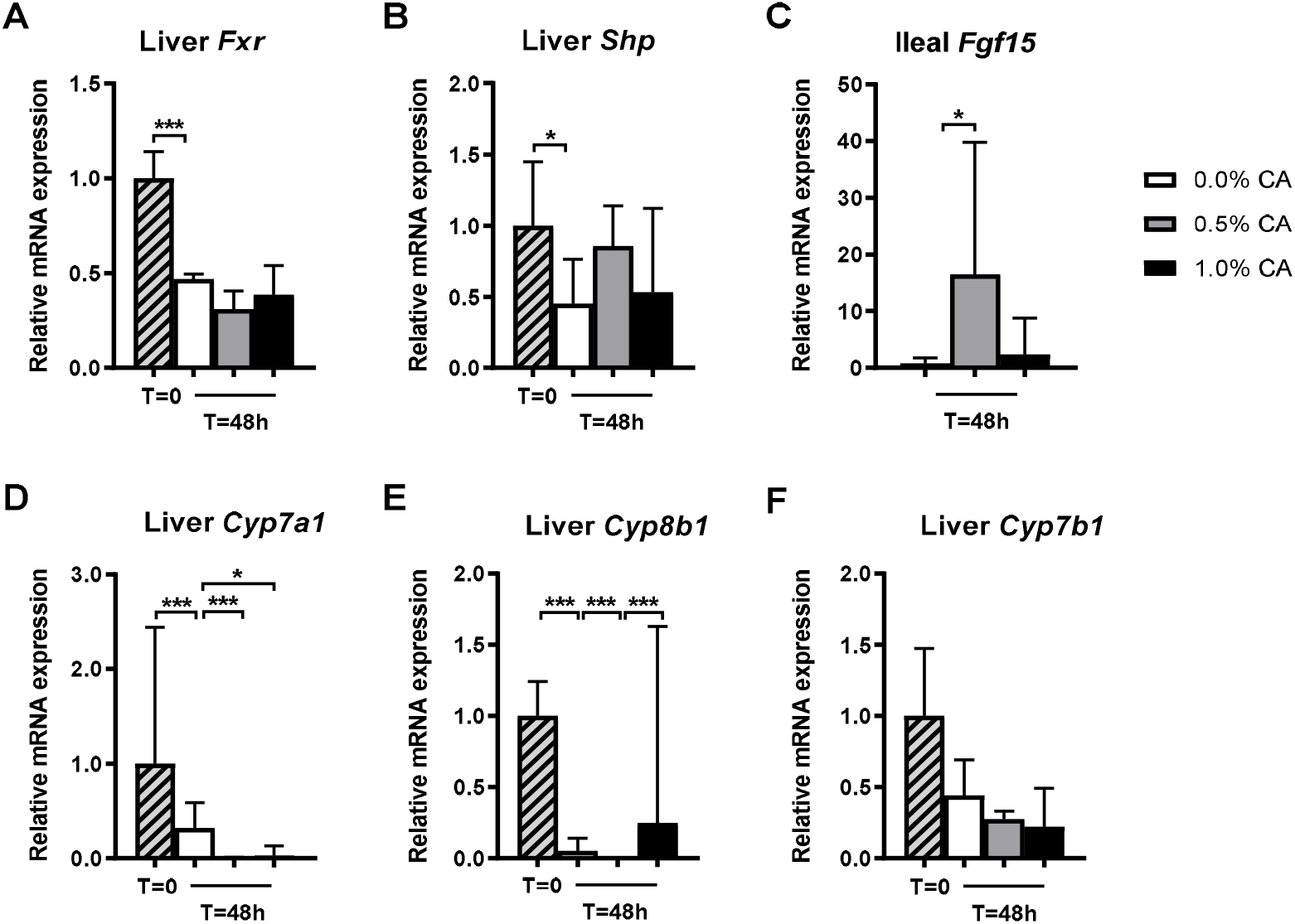
Suppression of bile salt synthetic genes in hepatectomized mice. Mice were subjected to 70% partial hepatectomy (PH) and post-operatively challenged with diet containing 0.0%, 0.5% or 1.0% cholic acid (CA, n=5-6 per group). Mice were sacrificed at 48 hrs after PH. Bile salt homeostasis was assessed after PH by activation of the Fxr-pathway in liver **(A**,**B)** and ileum **(C)**. *De novo* bile salt synthesis was assessed by expression of cytochrome P450 enzymes engaged in the classical **(D**,**E)** and acidic (**F**) synthetic pathways **(F)**. Values are expressed relative to the median expression in the control group. \**P*<0.05, \*\*\**P*<0.005. CA, cholic acid; Fxr, farnesoid X receptor; Shp, small heterodimer partner; Fgf, fibroblast growth factor; Cyp7a1, cholesterol 7α-hydroxylase; Cyp8b1, sterol 12α-hydroxylase; Cyp7b1, oxysterol 7α-hydroxylase.

In concordance with previous studies, PH induced a 4.1-fold reduction in *Cyp7a1* expression levels in mice fed a 0.0% CA diet (*P*<0.001; **Fig. 4D**), and further downregulation occurred in the 0.5% and 1.0% CA diet groups (*P*=0.004 resp. *P*=0.015). *Cyp8b1* expression levels followed the same pattern, although expression after PH was higher in mice fed 1.0% CA compared to those fed 0.5% CA (*P*=0.002) (**Fig. 5E**). *Cyp7b1* gene expression was neither influenced by PH nor by CA feeding (**Fig. 5F**).

**Figure 5.**
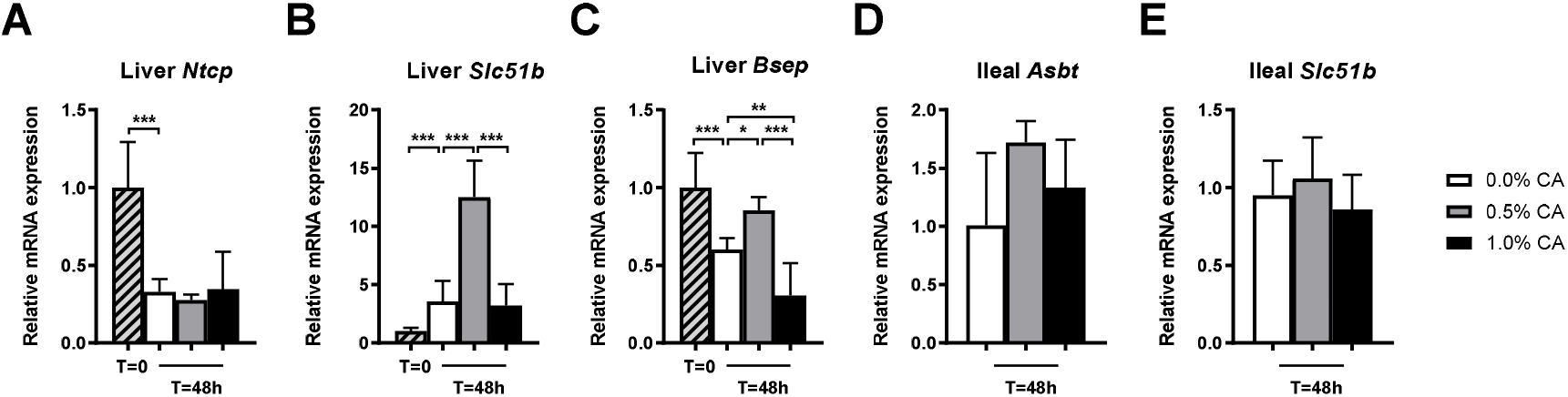
Expression of genes involved in hepatic and intestinal bile salt transport. Mice were subjected to 70% partial hepatectomy (PH) and post-operatively challenged with diet containing 0.0%, 0.5% or 1.0% cholic acid (CA, n=5-6 per group). Mice were sacrificed at 48 hrs after PH. Hepatocellular bile salt transport was assessed by gene expression of proteins engaged in uptake (**A**) and release (**B**,**C**) of bile salts. Ileal bile salt transport was assessed by gene expression of apical uptake (**D**) and basolateral export (**E**) transporters. Values are expressed relative to the median expression in the control group. \**P*<0.05, \*\**P*<0.01,\*\*\**P*<0.005. CA, cholic acid; Ntcp, sodium-taurocholate cotransporting polypeptide; Slc51b, organic solute transporter beta; Bsep, bile salt export pump; Asbt, apical sodium-dependent bile transporter.

#### 3.3.2 Effect of a post-operative cholic acid diet on bile salt uptake and export

Expression levels of genes engaged in the uptake and export of BS in liver and ileum were studied next. With regard to uptake transporters, a difference was only seen for hepatic *Ntcp* expression, namely a 3-fold reduction after PH (*P*<0.001), with no additional effects of CA feeding (**Fig. 5A**). PH caused marked upregulation (4-fold) of hepatic *Slc51b* expression (basolateral BS efflux, *P*<0.001) (**Fig. 5B**). This was further induced in mice fed a 0.5% CA diet, which had 3.1 resp. 3.9-fold higher *Slc51b* expression in relation to mice fed a 0.0% CA diet or a 1.0% CA diet (*P*=0.004 resp. *P*=0.002). PH induced a slight reduction (1.7-fold) in expression of *Bsep* (canalicular BS efflux, *P*=0.004), and feeding the mice a 0.5% CA diet partially restored expression (*P*=0.017 vs. 0.0% CA diet) (**Fig.5C**). In contrast, the 1.0% CA diet repressed *Bsep* expression relative to mice receiving a 0.0% or 0.5% CA diet (*P*=0.008 resp. *P*=0.002), and reduced entry of bile salts in the biliary tree and small intestine may underlie reduced ileal *Fgf15* expression in the 1.0% CA diet group. Along the same line, reduced canalicular (*Bsep*) and basolateral (*Slc51b*) bile salt secretion may contribute to elevation of liver injury markers **(Fig. 2AB)** in these mice.

No significant differences were seen in hepatic expression of transporters mediating canalicular secretion of bilirubin (*Mrp2*) and phospholipids (*Mdr3*), and basolateral secretion of BS (*Mrp3, Mrp4*) after PH (**Fig. S2**). In addition, CA feeding had no effect on expression of genes involved in intestinal uptake (*Asbt*) and secretion (*Slc51b)* of BS (**Fig. 6DE**).

**Figure 6.**
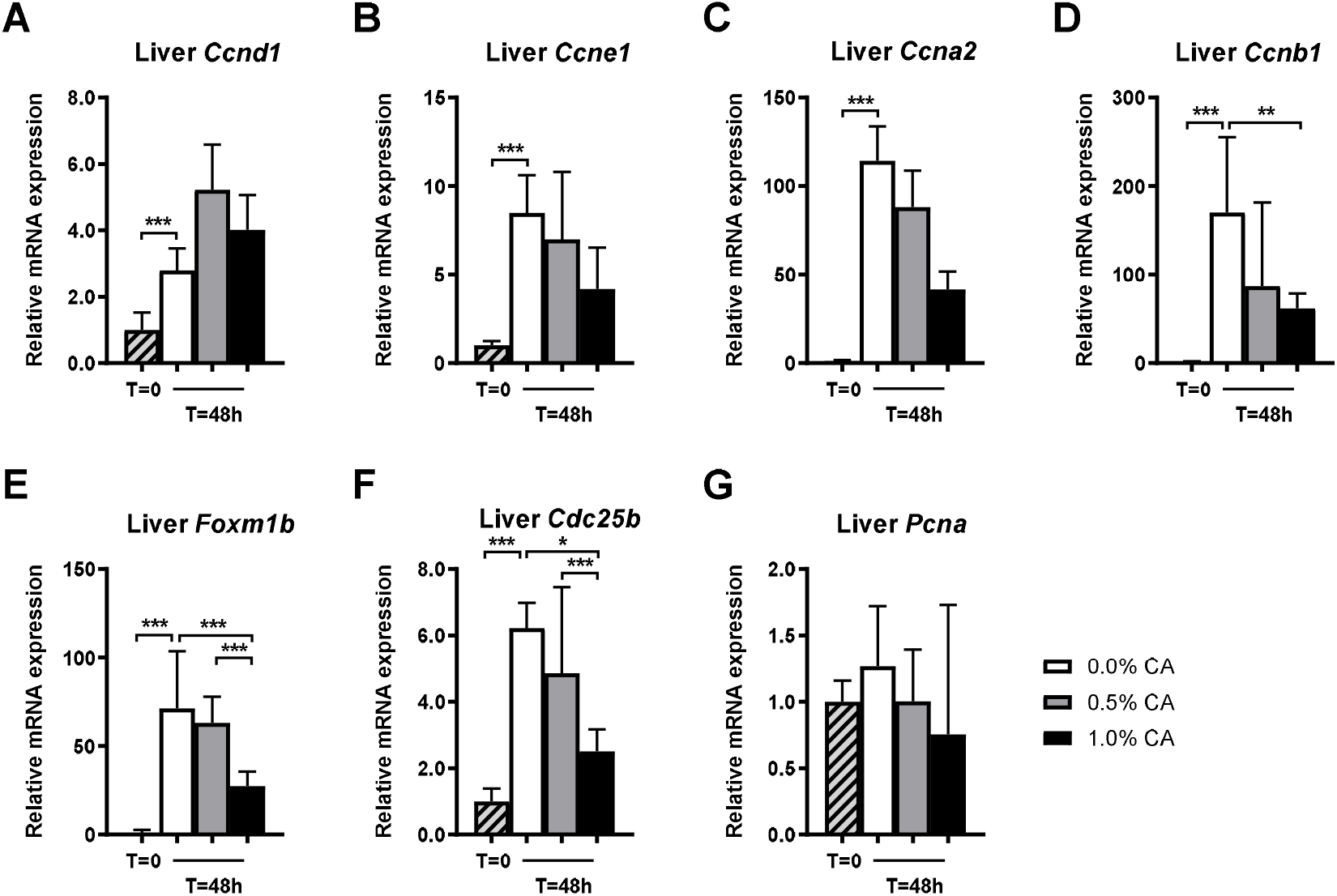
Abrogated cell cycle progression in hepatectomized mice fed a diet containing 1.0% cholic acid. Mice were subjected to 70% partial hepatectomy (PH) and post-operatively challenged with diet containing 0.0%, 0.5% or 1.0% cholic acid (CA, n=5-6 per group). Mice were sacrificed at 48 hrs after PH. Cell cycle progression was assessed by gene expression of enzymes involved in cell cycle regulation (**A-D**), entry into mitosis (**E, F**) and DNA synthesis (**G**). Values are expressed relative to the median expression in the control group. \**P*<0.05, \*\**P*<0.01,\*\*\**P*<0.005. CA, cholic acid; Ccn, cyclins; Fox, forkhead box; Cdc, cell division cycle; Pcna, proliferating cell nuclear antigen.

### 3.4 Effect of a post-operative cholic acid diet on cell cycle regulation and proliferation

Since hepatocyte proliferation was impaired in hepatectomized mice receiving a 1.0% CA diet, we examined hepatic expression of genes engaged in cell cycle regulation and proliferation. PH resulted in marked upregulation of cell cyclins *Ccnd1* (2.6-fold), *Ccne1* (8.8-fold), *Ccna2* (106-fold) and *Ccnb1* (159-fold) in control mice (*P*<0.002; **Fig. 6AD**), and with exception for *Ccnb1*, the extent of induction was comparable between diet groups. Mice fed a 1.0% CA diet had lower *Ccnb1* expression levels than mice fed a 0.0% CA diet (*P*=0.009).

With regard to expression of the cell cycle regulating transcription factor *Foxm1b*, whilst expression was virtually undetectable in quiescent liver, PH caused a marked induction (64-fold) (*P*<0.001; **Fig. 6E**). When comparing the different dose groups, the 1.0% CA group showed abrogated PH-induced *Foxm1b* expression compared to both the 0.0% and 0.5% CA diet groups (both *P*=0.004). Gene expression level of cell division cycle 25b (*Cdc25b*), a direct target gene of *Foxm1b* and required for entry into mitosis, was induced by PH (*P*<0.001) (**Fig. 6F**). Analogous to *Foxm1b*, the 1.0% CA group displayed an impaired induction of *Cdc25b* compared to mice receiving a 0.0% or 0.5% CA diet (*P*=0.017 resp. *P*=0.002). PH or CA feeding had no effect on expression of proliferating cell nuclear antigen (*Pcna*) (**Fig. 6G**).

## 4. Discussion

PLF is a serious complication following liver resection with high morbidity and mortality. In order to find an appropriate therapy to treat and/or prevent the occurrence of liver failure after resection, an animal model of PLF is of great value. We hypothesized that excessive accumulation of BS in the regenerating liver remnant is the actual culprit in PLF. To test this, we induced BS overload in the regenerating liver of mice by feeding them a CA diet after 70% PH. Concentrations of CA in the diet ranged from 0.0 to 1.0%, and mice were sacrificed around the time of maximal hepatocyte proliferation (normally peaking between 36-48 hrs).

Mice in the highest dose group had poorer ‘clinical’ performance (*i*.*e*. squinted eyes, reduced physical activity) as indicated by our welfare assessment (**Fig. S1**), with overall decreased glucose levels after PH, and more pronounced body weight loss in the postresectional course (**Fig. 1AB**). Moreover, in this group, assessment of markers for liver injury (aminotransferases) and secretory function (total bilirubin) revealed hepatocellular injury and an impaired secretory function (**Fig. 2**). Although no effect of a 1.0 % CA diet was seen on liver mass recovery, impaired hepatocyte proliferation was noted (**Fig. 1CD**).

In contrast, despite a cholic acid-supplemented diet, mice fed a 0.5% CA diet seemed to perform similar or even better than the 0.0% CA diet group in terms of hepatocyte proliferation. Ileal *Fgf15* was upregulated in mice fed a 0.5% CA diet compared to a 0.0% CA diet (**Fig. 4C**). No signs of liver injury and a maintained hepatocytic proliferative capacity were seen in the 0.5% CA group. Since FGF19/Fgf15 has been identified as a direct hepatic mitogen, maintained proliferative capacity in the 0.5% CA group may relate to enhanced ileal *Fgf15* expression [8].

In our study, a reduction in hepatic BS content was seen in control mice 48 hrs after PH (**Fig. 3A**). We studied gene expression of Fxr-regulated genes to find a mechanistic explanation for this. PH is known to result in a relative overload of BS in the remnant liver directly after PH, likely resulting in increased bile salt signaling via Fxr and induction of Fxr target genes, such as *Shp* and *Slc51b* [14]. Previously, Uriarte et al. demonstrated a transient increase in intrahepatic BS levels 24 hours after PH, followed by a decrease to values almost equal to baseline levels after 48 hours [8]. This is in accordance with Huang et al., who showed that PH induced a decrease in hepatic BS content in mice after 48 hours [15]. The decreased hepatic BS content 48 hours after PH may be linked to Fxr/Fgf15-mediated repression of bile salt synthesis (*Cyp7a1*) and downregulation of the hepatocytic BS uptake transporter *Ntcp* in reaction to the relative overload [5], which was also observed in our study (**Fig. 4D,5A**).

Although an elevation in hepatic BS content was seen in both CA-fed groups compared to a 0.0% CA diet, no difference was found between the various dose groups. These results were not in accordance with the expectation that a 1.0% CA diet would result in more accumulation of BS in the remnant liver and consequent impaired liver mass regrowth. Possible differences in hepatic bile salt composition, an important determinant of cytotoxicity, have not been assessed. Likewise, hepatic BS measurements are inevitably based on homogenates, and information on the actual spatial localization (i.e. within the hepatocytes, within the biliary network) of BS in the context of liver regeneration is not available. Yet, it is plausible that in particular accumulation of BS within the hepatocytes, negatively impacts hepatocyte proliferation. Divergent changes in *Bsep* expression in the CA diet groups, favoring localization of BS within the biliary network in the group (0.5% CA diet) that had highest percentage of proliferating hepatocytes, supports this idea. To determine the spatial distribution of distinct bile salts species in liver sections and thereby investigate local actions, techniques such as matrix-assisted laser desorption/ionization-mass spectrometry imaging (MALDI-MSI) have proven profitable [16]. CA feeding led to an elevation of serum BS in the 0.5% CA group only. This can be interpreted as adaptive mechanisms that serve to maintain low intracellular BS levels, e.g. enhanced basolateral (via Slc51b) and canalicular export (via Bsep), being functional in the 0.5% CA group. By extension, these protective mechanisms may have failed in the 1.0% CA diet group. Again, data on spatial localization of BS in the CA groups, would be informative.

Postresectional hepatocellular proliferation proceeds through tightly regulated transitions which are, amongst others, controlled by cell cycle regulatory genes. PH induced an upregulation in gene expression of all cyclins that were studied, as well as Foxm1b and Cdc25b. This is in accordance with the literature, and the consequence of cell cycle re-entry upon hepatic resection [17]. Importantly, mice fed a 1.0% CA diet had lower PH-induced Ccnb1, Foxm1b and Cdc25b levels compared to mice fed a 0.0% or 0.5% CA diet (**Fig. 6**), and this translated to the reduced percentage of proliferating (*i*.*e*. Ki-67^+^) hepatocytes (**Fig. 1D**). This means that cell cycle progression is hampered in the 1.0% CA group. Impaired cell proliferation could be due to initiation of liver injury that interferes with proper regeneration. In contrast with the study of Uriarte et al., we did not find any differences in Pcna at the mRNA level (**Fig. 6G**).

After PH, liver mass recovery is often used as a regeneration index [18,19]. It is however a rough estimate based on the wet weight that assumes removal of exactly 70% of the liver in all cases, and does not consider unrelated causes of liver mass gain although phenomena such as steatosis and hepatocyte hypertrophy are associated with regeneration. Especially with regard to the poor clinical performance of mice fed a 1.0% CA diet, for instance hepatic edema may have affected wet liver weight measurement. In this regard, direct indices to assess hyperplasia of the liver, such as Ki-67 staining that was employed here, may be more informative.

In conclusion, a postresectional BS challenge of 1.0% CA induced signs of liver injury and resulted in impaired liver regeneration. Based on these data, we can conclude that the approach (postresectional CA-challenge) to establish a PLF model looks promising, but modifications to the protocol are required. To test if the pre-damaged liver is extra susceptible for (bile salt) injury, we could apply the current model in mice with cholestatic liver disease due to occlusion of the common bile duct.

## Abbreviations

PLF: postresectional liver failure
BS: bile salts;
NR: nuclear receptors;
FXR: farnesoid X receptor;
PXR: pregnane X receptor;
BSEP: bile salt export pump;
FGF: fibroblast growth factor;
SHP: small heterodimer partner;
ALF: acute liver failure;
PH: partial hepatectomy;
CA: cholic acid;
ALT: alanine aminotransferase;
AST: aspartate aminotransferase

## Supplemental data

**Supplementary Figure 1.**
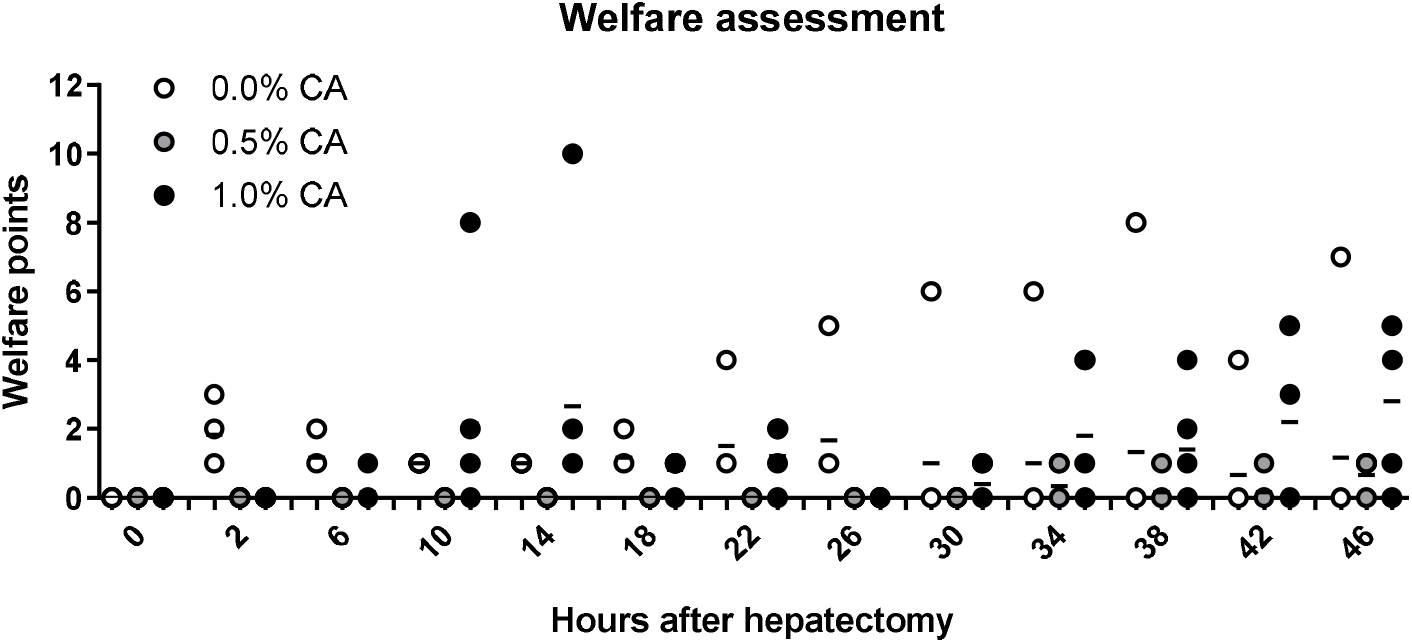
Increased welfare scores in mice fed a 1.0% cholic acid diet. Mice were subjected to 70% partial hepatectomy (PH) and post-operatively challenged with diet containing 0.0%, 0.5% or 1.0% cholic acid (CA, n=5-6 per group). Mice were sacrificed at 48 hrs after PH. Welfare was assessed every 4 hrs from 2 hrs after PH onwards, by a specific score list (Table S1). A higher total of welfare points indicates a lower welfare. Data are presented as individual points with median (aligned scatter plot). *mouse excluded from analysis due to technical failure of PH.

**Supplementary Figure 2.**
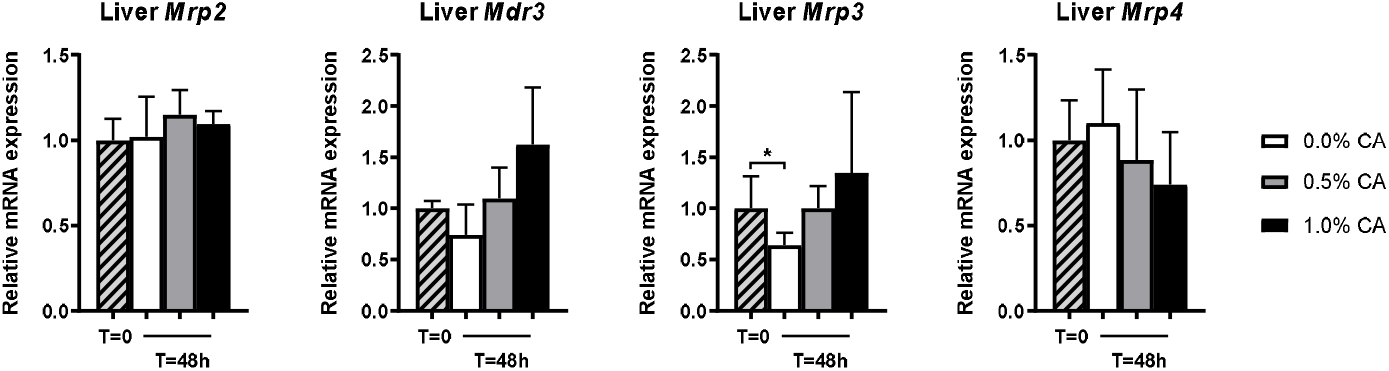
Unaltered gene expression of proteins engaged in basolateral and canalicular secretion of cholephiles in mice fed a diet containing 1.0% cholic acid. Mice were subjected to 70% partial hepatectomy (PH) and post-operatively challenged with diet containing 0.0%, 0.5% or 1.0% cholic acid (CA, n=5-6 per group). Mice were sacrificed at 48 hrs after liver resection. Canalicular secretion of bilirubin **(A)** and phospholipids **(B)**, and basolateral secretion of BS **(C**,**D)** was assessed by analyzing gene expression of relevant transporters. Data are presented as median with interquartile range. \**P*<0.05. CA, cholic acid; Mrp, multidrug resistance-related protein; Mdr, multidrug resistance protein.

**Supplementary Table 1.**
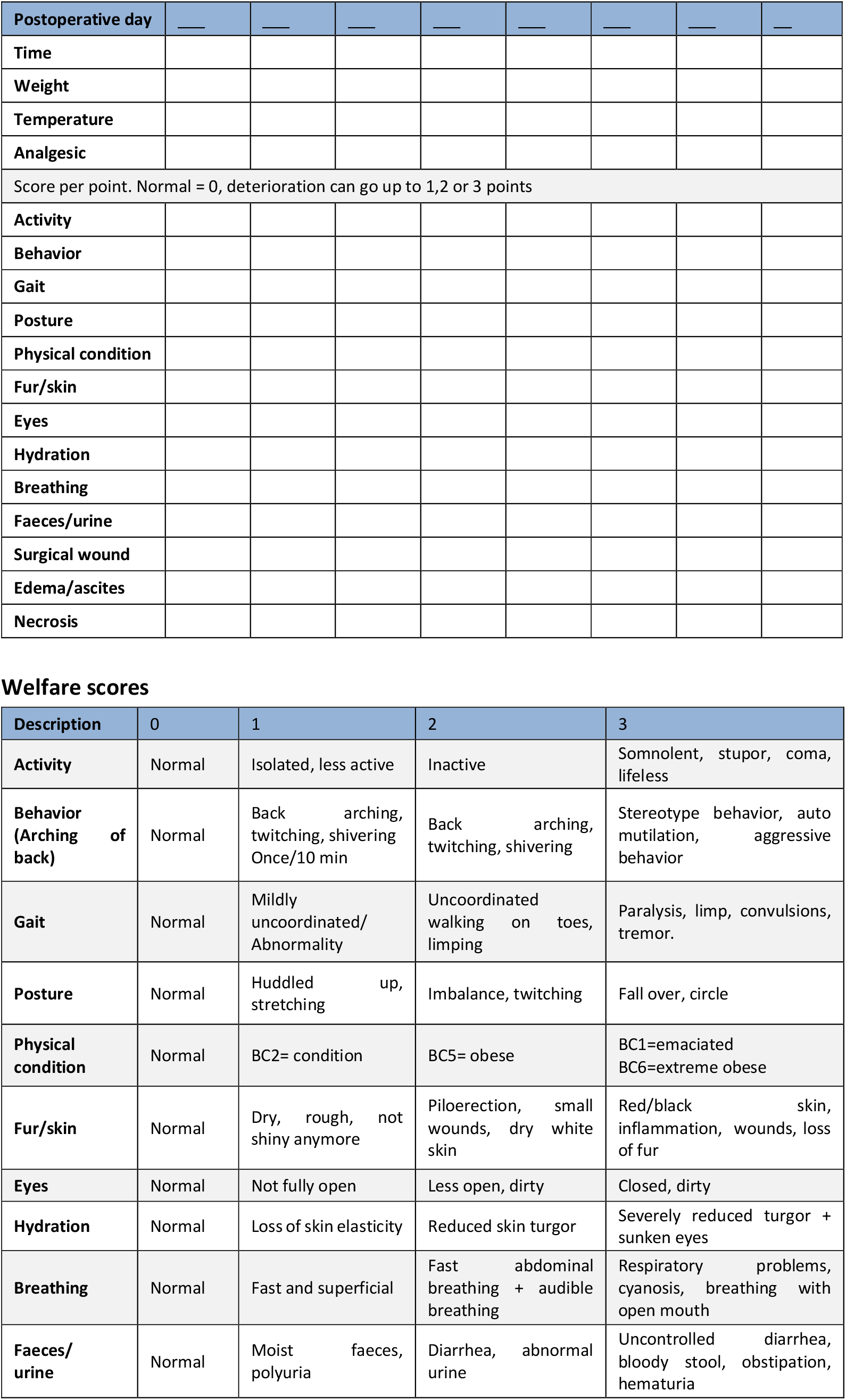
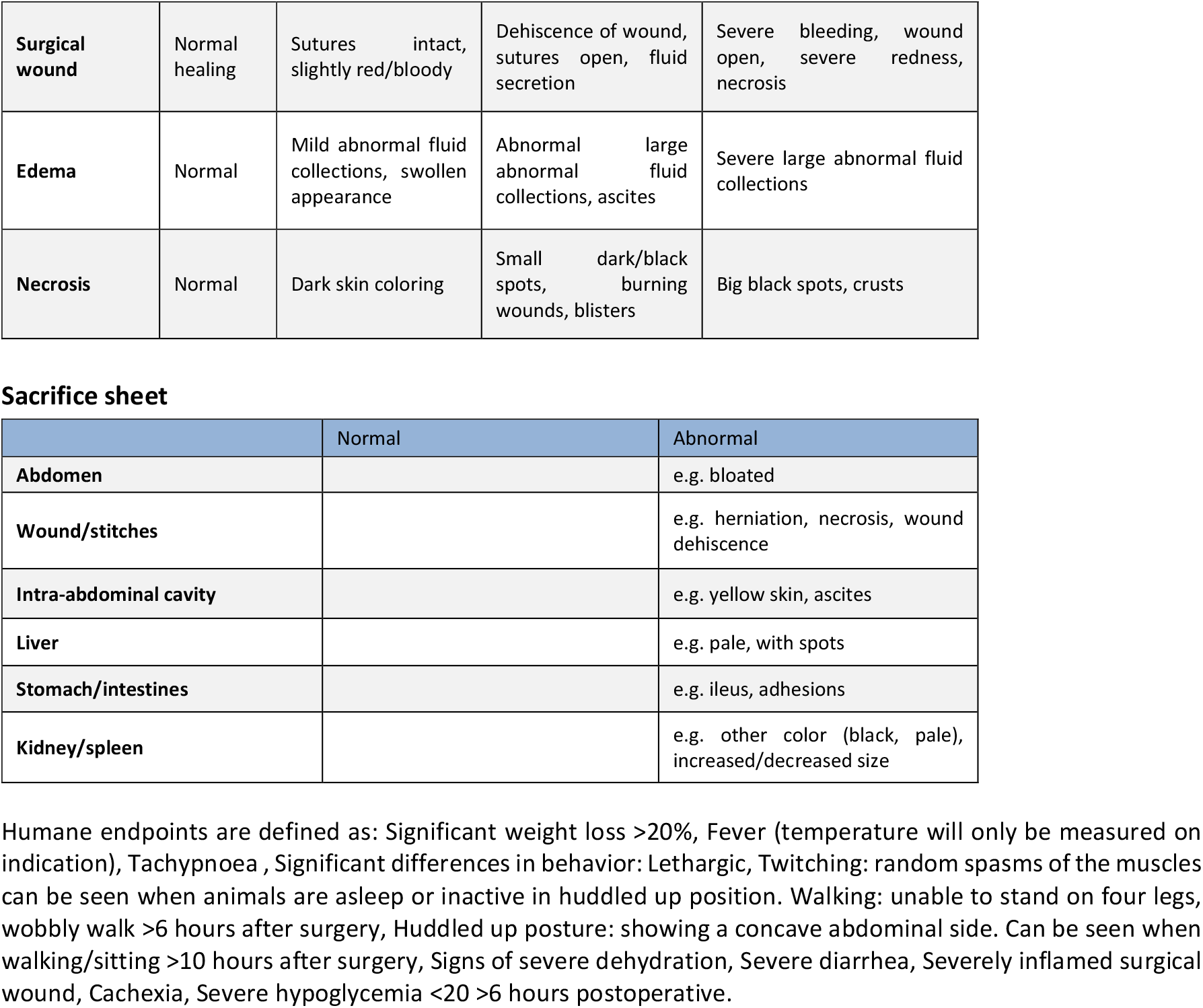
Postoperative welfare assessment form.

**Supplementary Table 2.**
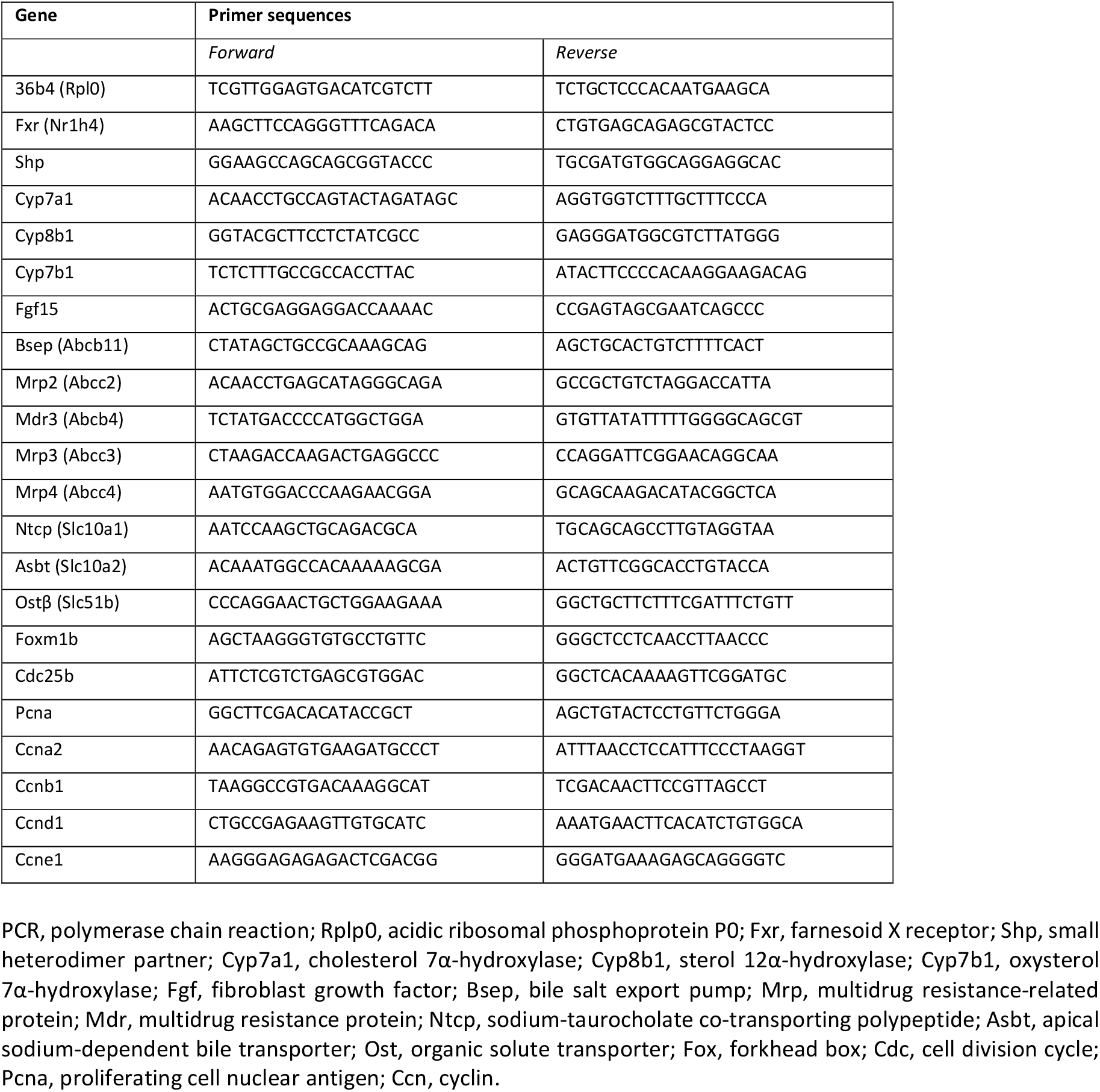
Primer sequences used in RT-qPCR.

